# Relating dynamic brain states to dynamic machine states: human and machine solutions to the speech recognition problem

**DOI:** 10.1101/074799

**Authors:** Cai Wingfield, Li Su, Xunying Liu, Chao Zhang, Phil Woodland, Andrew Thwaites, Elisabeth Fonteneau, William D Marslen-Wilson

## Abstract

There is widespread interest in the relationship between the neurobiological systems supporting human cognition and emerging computational systems capable of emulating these capacities. Human speech comprehension, poorly understood as a neurobiological process, is an important case in point. Automatic Speech Recognition (ASR) systems with near-human levels of performance are now available, which provide a computationally explicit solution for the recognition of words in continuous speech. This research aims to bridge the gap between speech recognition processes in humans and machines, using novel multivariate techniques to compare incremental 'machine states', generated as the ASR analysis progresses over time, to the incremental 'brain states', measured using combined electro-and magneto-encephalography (EMEG), generated as the same inputs are heard by human listeners. This direct comparison of dynamic human and machine internal states, as they respond to the same incrementally delivered sensory input, revealed a significant correspondence between neural response patterns in human superior temporal cortex and the structural properties of ASR-derived phonetic models. Spatially coherent patches in human temporal cortex responded selectively to individual phonetic features defined on the basis of machine-extracted regularities in the speech to lexicon mapping process. These results demonstrate the feasibility of relating human and ASR solutions to the problem of speech recognition, and suggest the potential for further studies relating complex neural computations in human speech comprehension to the rapidly evolving ASR systems that address the same problem domain.

**Author Summary:** The ability to understand spoken language is a defining human capacity. But despite decades of research, there is still no well-specified account of how sound entering the ear is neurally interpreted as a sequence of meaningful words. At the same time, modern computer-based Automatic Speech Recognition (ASR) systems are capable of near-human levels of performance, especially where word-identification is concerned. In this research we aim to bridge the gap between human and machine solutions to speech recognition. We use a novel combination of neuroimaging and statistical methods to relate human and machine internal states that are dynamically generated as spoken words are heard by human listeners and analysed by ASR systems. We find that the stable regularities discovered by the ASR process, linking speech input to phonetic labels, can be significantly related to the regularities extracted in the human brain. Both systems may have in common a representation of these regularities in terms of articulatory phonetic features, consistent with an analysis process which recovers the articulatory gestures that generated the speech. These results suggest a possible partnership between human-and machine-based research which may deliver both a better understanding of how the human brain provides such a robust solution to speech understanding, and generate insights that enhance the performance of future ASR systems.

## 1 Introduction

A fundamental concern in the human sciences is to relate the study of the neurobiological systems supporting complex human cognitive functions to the development of computational systems capable of emulating or even surpassing these capacities. Spoken language comprehension is a salient domain that depends on the capacity to recognise fluent speech, decoding word identities and their meanings from a stream of rapidly varying auditory input.

In humans, these capacities depend on a highly dynamic set of electrophysiological processes in speech-and language-related brain areas. These processes extract salient phonetic cues which are mapped onto abstract word identities as a basis for linguistic interpretation. But the exact nature of these processes, their computational content, and the organisation of the neural systems that support them, are far from being understood. The rapid, parallel development of Automatic Speech Recognition (ASR) systems, with near-human levels of performance, means that computationally specific solutions to the speech recognition problem are now emerging, built primarily for the goal of optimising accuracy, with little reference to potential neurobiological constraints.

In the research described here we ask whether the properties of such a machine solution, in terms of the input-output relations it encodes, can be used to probe the properties of the human solution, both to develop new insights into the properties of human speech analysis and to suggest new constraints on the analysis strategies of future ASR systems. We do not seek to invoke the specific properties of the computational architectures of machine systems and of human systems (see Section 3.2), but rather to ask, in this initial study, whether the regularities that successful ASR systems find between information in the speech input and word-level phonetic labelling can be related to the regularities extracted by the human system as it processes and identifies parallel sets of words.

A critical issue in doing so is to capture the incremental and temporally distributed nature of the speech input and its interpretation, where partial cues to phonetic and lexical interpretation build up gradually over periods of potentially hundreds of milliseconds (in contrast to visual perception of static objects), so that processes at all levels of description are continuously modified and overlaid as new sensory constraints emerge over time. For this study, therefore, we need to define incremental ‘machine states’, capturing the multi-level labelling assigned to a given speech input as the ASR analysis progresses over time, and relate these to incremental ‘brain states’ generated by human listeners as the same inputs are heard and perceived.

### 1.1 Multivariate methods for probing real-time brain states

The possibility of engaging on this enterprise depends on new methods for non-invasively investigating the real-time electrophysiological activity of the human brain. We build here on earlier work [1] which used a novel multivariate pattern analysis method, called Spatiotemporal Searchlight Representational Similarity Analysis (ssRSA), to decode information about frequency preference and selectivity directly from the dynamic neural activity of the brain as reconstructed in combined MEG and EEG (EMEG) source space. This method is an extension of fMRI-based RSA [2, 3] to time-resolved imaging modalities.

The core procedure in ssRSA is the computation of similarity structures that express the dynamic patterns of neural activation at specific points in space and time. This similarity structure is encoded in a representational dissimilarity matrix (RDM), where each cell in the RDM records a measure of the dissimilarity between the neural activation patterns elicited by pairs of experimental conditions (for example, sets of auditory stimuli). These brain data RDMs, reflecting the pattern of brain activity within the spatiotemporal window defined by the searchlight procedure [2, 4, 5], are then related to model RDMs, which express specific theoretical hypotheses about the properties of this activity. In our previous study [1], exploring the frequency preferences and selectivity of auditory cortex, the model RDMs capture the similarity between each stimulus at each frequency band, derived from a computational model of the early stages of auditory processing [6]. The ssRSA procedure made it possible to relate low-level neural patterns directly to abstract higher-level functional hypotheses about the organization of the auditory cortex.

In the current research we use ssRSA to compute representations of the similarity structure of the brain states generated incrementally as human listeners perceive and interpret spoken words. These brain data RDMs can then be related to time-varying model RDMs which capture the similarity structure of the machine states extracted during an ASR analysis of the same sets of spoken words. Critically, since these comparisons were conducted in terms of the abstract geometric configuration [7] of the mappings captured in the data RDMs and the model RDMs, this allows us to compare the properties of machine solutions and human solutions to the recognition problem without assuming any correspondence between the specific format of the state information in the ASR system, and the expression of information represented in the human system — or, indeed, between the functional architectures of the two systems.

### 1.2 Shared representational frameworks: Phones and features

The historically dominant approach to speech recognition, whether in human or machine contexts, assumes that the mapping from speech input to linguistic interpretation is mediated through stored form representations of the words in the language, and that access to these representations is in terms of a phonetic labelling of the speech input (though see [8]). This labelling is generated as the speech input is incrementally analysed over time. This potentially provides the basis for a shared *representational framework* for relating the content of machine solutions on the one hand to assumed human solutions on the other. As the ASR system computes a probability distribution over a number of discrete phonetic labels for the speech input, this can form the basis for RSA model RDMs which in turn can be related to human brain data RDMs, where these are thought to reflect the computation of similar types of regularity in the speech stream.

This leaves open, however, the question of what is the most appropriate form for this shared representational framework. Here we proceed on the assumption that the human solution is best captured in terms of articulatory phonetic features rather than in terms of classic phonetic or phonemic labels. These features seek to capture the underlying articulatory gestures by which the human speech output is generated, and represent a neurobiologically plausible, ‘low-dimensional’ solution to the speech analysis problem. Such feature sets have a long history in linguistics and in the speech sciences, while substantial behavioural evidence supports their role in the incremental speech analysis process (e.g., [9, 10]). In particular, feature-based accounts allow more natural treatment of the partial acoustic-phonetic information that becomes available as overlapping articulatory gestures incrementally generate the speech output. This is also a point that has been long made by some ASR researchers (e.g. [11]). There is also substantial evidence from neuroimaging studies that the neural substrates for speech processing can be characterised in featural terms. Several studies by Obleser and colleagues [12–14], for example, using both fMRI and MEG, show that neural responses distributed across auditory cortices in superior temporal regions reliably differentiate between phonetic features associated with both vowels and consonants (see also [15]). Specific evidence for the role of articulatory features comes in particular from studies in the “motor theory” tradition, which claims that the perception of speech is directly structured by the recovery of the articulatory gestures that generated the speech output being heard (e.g., [16–18]. Most directly relevant, however, in the neurophysiological context of the current project, is the recent research using electrocorticographic (ECoG) techniques where intracranial electrodes are used to record directly from bilateral speech processing areas in human superior temporal cortex [19]. This work shows that the patterns of neural response to spoken sentences correspond well to an analysis in terms of articulatory features. Taking individual ECoG electrode responses as feature vectors for each phone, Mesgarani et al. [19] showed that the phones cluster together in a manner well-described by a set of articulatory features. While this research is not able to definitively pull apart phone-based from feature-based approaches, it confirms the viability of featural decomposition as a means of characterising earlier stages of speech analysis in the human brain.

Here we use a standard set of such features, based on those used by Mesgarani et al. [19], but adapted for British rather than American English [20, 21]. Most ASR systems, in contrast, have chosen phones (the inventory of distinct speech sounds in a given language) rather than features as their intermediate representational unit. This means that, analogous to the procedure used by Mesgarani et al. [19], we can use the ASR system to generate incremental streams of probabilistic phonetic labels, and then assess these as evidence for the presence of the corresponding underlying articulatory features. The hidden Markov model toolkit (HTK) Version 3.4 [22], the ASR system we use here, is based on a successful GMM-HMM architecture (with word-level accuracy of around 91.6%). It uses a Gaussian mixture model (GMM) to annotate each successive 10 ms frame of a recorded speech stream with the estimated likelihood of each phonetic label and each triphone label, while a hidden Markov model (HMM) captures statistical regularities in the language and its sequential structure. The triphones capture the phonetic context of each frame, enabling the ASR system to take into account important co-articulatory variation in the phonetic form of a given phone, and comprise a triplet indicating the current phone, as well as a preceding and future phone. For example, [p-i+n] is a [i] as pronounced with the preceding context of a [p] and the following context of a [n]. As we describe below in Section 2.2, the triphone likelihoods generated at each 10 ms time-step (over a 60 ms time-window) for each speech stimulus can be used to generate RSA model RDMs for each phone in the language. The resulting 40 phonetic model RDMs are then tested against the corresponding brain data RDMs in a searchlight procedure across relevant brain areas. The resulting distributions of phonetic model fits at each testing point are then used to derive an account in terms of phonetic features.

### 1.3 Spatiotemporal foci of the analyses

The brain data used in this study come from simultaneously recorded EEG and MEG (EMEG) measurements of the neural activity generated by each of 400 spoken words heard by the listeners. Critically, we use these EMEG data, as in the earlier Su et al. [1] study, to generate a ‘source space’ reconstruction that localises at the cortical surface (specifically, the white matter/grey matter boundary) the electrophysiological activity that gives rise to the EMEG measurements recorded at sensors external to the skull. The combination of MEG and EEG delivers better source localisation than either of these modalities alone [23], using well established minimum norm estimation (MNE) techniques guided by neuroanatomical constraints from structural MR for each participant [24, 25].

These source space representations allow us to track, with millisecond temporal resolution and potentially sub-centimetre spatial resolution, the real-time spatiotemporal patterns of electrophysiological activity generated by the brain as it performs complex operations such as speech interpretation. This means that we can focus our explorations of this neural activity on those regions of cortex which are known to play a key role in supporting the operations of interest rather than, for example, working with the pattern of MEG data measured in ‘sensor space’ (e.g. as in [26], where sources are not localised). In the current exploration, therefore, of the potential relationship between the incremental brain states derived from the source-localised EMEG data and the incremental machine states derived from ASR responses to the same 400 spoken words, we restrict the scope of these analyses to the spatiotemporal locations in the brain data where such a relationship is most likely to be found.

In terms of spatial locations, as demonstrated in recent ECoG studies, (e.g. [19, 27]), as well as in a host of neuroimaging experiments using fMRI techniques, the superior temporal cortices bilaterally, including Heschl’s gyrus, are the key areas supporting acoustic-phonetic analysis and the earlier stages of speech interpretation (e.g. [28, 29]). Accordingly, we restrict the analyses reported here to a *speech area mask* covering superior temporal cortex (STC) bilaterally.

The second critical dimension concerns the timing of the processing lag (in milliseconds) between the arrival of information in the speech input and the acoustic-phonetic neural interpretation of this information in superior temporal cortex. This lag reflects both neural transmission time (from cochlea to cortex), and the processing time necessary to compute the phonetic distinctions reflected in the human cortical response. Neither of these sources of delay apply to the representations computed by the ASR system. This means that a compensatory lag must be introduced into the matching between the ASR-based model RDMs for a given time-window of speech input and the data RDMs representing the human listeners’ cortical response to this same stretch of input.

Evidence for the timing of such a lag has become available from recent research relating speech input to higher-order cortical interpretation. In a study using EMEG word data similar to the speech materials used here, Thwaites et al. [30] found that the cortical computation of the fundamental frequency function *F*0 was most strongly expressed in EMEG brain responses in bilateral STC at a lag of 90 ms. Related evidence from ECoG studies looking at phonetic discriminability in neural responses to speech suggest similar delays. Mesgarani et al. ([19], Fig S4A) show a lag of around 125 ms for neural discrimination of phonetic categories relative to the acoustic information in the speech signal, while Chang et al. ([27], Fig 2a) find a similar peak at around 110 ms. Accordingly in our analyses here we use a fixed lag of 100 ms, so that each time-window of ASR-derived model RDM data is centred over the time-window of brain data RDMs recorded (from locations in STC) 100 ms later. In later studies we hope to be able to test at a wider range of lags.

### 1.4 Overview

Our primary goals in this project are to investigate the basic technical and conceptual feasibility of relating dynamic brain states to dynamic machine states, in the incremental processing environment imposed by the speech input, and to determine whether substantive correspondences can indeed be found between human and ASR solutions to the speech recognition problems. In so doing, we will ask whether a potential account of these commonalities in terms of articulatory phonetic features proves to be viable. In the following sections of the paper we describe the sets of procedures and analyses required to achieve these goals.

Section 2.1 provides an overview of the RSA approach in the EMEG context. Section 2.2 covers the RSA method for computing the ASR phone model RDMs from the incremental output of HTK. In Section 2.3 we report a novel subsidiary analysis, conducted to evaluate the applicability of a specific articulatory feature set to the resulting 40 phone-specific dynamic model RDMs. We then turn (Section 2.4) to the procedures for computing data RDMs from incremental brain states, with two subsections (2.4.1 and 2.4.2) covering source reconstruction and the RSA searchlight process. Section 2.5 lays out the core multivariate RSA process of (a) fitting the machine model RDMs to the brain data RDMs, conducted in superior temporal cortex at the chosen 100 ms lag, and (b) converting these phone model data to an account in terms of articulatory features. The relevant permutation-based statistical apparatus is outlined in Section 2.5.1, while 2.6 presents the outcome of the RSA analyses that relate computational models of speech analysis extracted from ASR machine states to the neural speech analysis processes non-invasively detectable in human superior temporal cortex.

## 2 Procedures and Analyses

### 2.1 Representational similarity analysis for EMEG

RSA was originally developed for fMRI [5], but one of its strengths is its agnosticism toward modality. In particular, RSA has more recently been applied to EEG and MEG data [1, 31]. Methodological considerations for EMEG are detailed in Su et al.’s work [4], but we will give a brief overview here.

RSA involves the comparison of the similarity (or dissimilarity) structures found in the responses of neural systems to differing experimental conditions, and the modelling of those similarity structures by categorical or computational models. A basic component of the RSA analysis strategy is the construction of representational dissimilarity matrices (RDMs) based on the pairwise dissimilarity of all items or conditions entered into the analysis process [1, 5]. Typically, the data RDM or set of data RDMs is computed from a neuroimaging data source, and the model RDM(s) from a specified model of the stimuli. A generic description of this process is provided here and illustrated schematically in Fig 1, while the specific choices and parameters used in this analysis are explained in Section 2.4.

**Figure 1:**
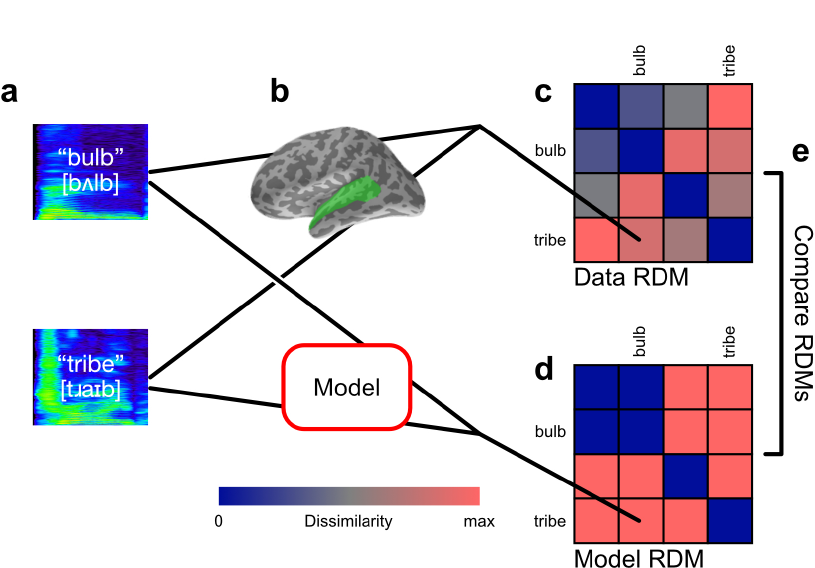
Representational similarity analysis. **(a)** A set of experimental conditions or stimuli are presented to participants. In this example, recordings of English words are presented aurally. **(b)** For each experimental condition, EMEG data is collected from participants’ regions of interest for a specified epoch. **(c)** Dissimilarities between each pair of responses are computed and stored in a representational dissimilarity matrix. Potential dissimilarity measures include Pearson’s correlation distance or Euclidean distance between response vectors. Rows and columns of the matrix are indexed by the condition labels, making the matrix symmetric with diagonal entries all 0 by definition. In this example there are four conditions in total, and the responses to the condition pair *(bulb, tribe*) is compared, with the value stored in the indicated matrix entry, and its diagonally-symmetric counterpart. **(d)** A model of the experimental conditions or stimuli is used to compute a model RDM. The model RDM can be computed in several ways, e.g. by comparing representations of the stimuli under the model; or by modelling the dissimilarities directly. **(e)** Data and model RDMs are statistically compared, e.g. by computing Spearman’s rank correlation of their upper-triangular vectors.

For EMEG data, field strength is measured at each of the sensor sites in the MEG helmet and at the EEG scalp electrodes as the participant is presented with an experimental trial, such as an auditory stimulus (Fig 1a-b). The dissimilarities between the responses to each pair of stimuli are computed using some distance measure, often Pearson’s correlation distance between the vectors of measured response. These distances are entered into the data RDM (Fig 1c). Data RDMs are computed for each subject of the experiment, and may be averaged between subjects [5]. The particular choices made in this study are detailed in Section 2.4.

The model RDM is an analogous matrix of predicted dissimilarities between experimental conditions (Fig 1d). It may be produced in several ways, such as a pairwise comparison of model variables for each stimuli, or by directly modelling the dissimilarity values themselves.

Because the rows and columns of the data RDM and model RDM are both indexed by the experimental conditions, they can be directly compared (Fig 1e). This comparison often takes the form of a correlation of their upper-triangular vectors, yielding a correlation value describing how well the dissimilarities expressed in the model RDM explain the dissimilarities in the data RDM. A rank correlation such as Spearman’s may be used rather than a linear correlation such as Pearson’s, so that a strictly linear relationship between modelled and actual dissimilarities is not predicted.

### 2.2 Computing model RDMs from incremental machine states

For this study, model RDMs were computed based on dissimilarities between modelled representations of stimuli, rather than directly modelling the dissimilarities themselves. The models were constrained to the set of English words for which we had suitable EMEG brain data. These were 400 English verbs and nouns (e.g., *talk, claim*) some of which had past tense inflections (e.g., *arrived, jumped*). These materials (also used in [1]) were prepared for another experiment [32], and we assume (a) that their linguistic properties were independent of the basic acoustic-phonetic parameters being examined here and (b) that they provide a reasonable preliminary sample of naturally occurring phonetic variants in human speech. The stimuli were recorded in a sound-attenuated room by a female native speaker of British English onto a DAT recorder, digitized at a sampling rate of 22 kHz with 16-bit conversion, and stored as separate files using Adobe Audition (Adobe Inc., San Jose, CA). They averaged 593 ms in length, with the shortest word being 270 ms in length. To keep the speech samples and analyses fully aligned we restricted all the analyses to the first 270 ms of each word.

A hidden Markov model (HMM) based automatic speech recognition (ASR) system with Gaussian mixture model (GMM) observation density function was built using HTK [22]. The acoustic training corpus used to build the ASR system was the 15 hours Wall Street Journal corpus with British accents recorded at Cambridge University (WSJ-CAM0). There are overall 44 phones defined in the corpus, which can result in a maximum number of 85,184 triphone units. Following a well-established strategy [22], each HMM used 3 hidden states to model a single triphone unit. To have sufficient data for a more reliable estimate of all HMM model parameters and to model the triphone units that did not occur in the training set, the standard decision tree based tying approach was adopted to cluster the hidden states with the same centre phone unit according to phonetic knowledge and a maximum log-likelihood criterion [33].

After building the ASR system, the 400 recorded stimulus words were recognized with the GMM-HMM models, with a modified version of the HTK recognizer that output the triphone likelihoods that it generated as it processed the stimuli (in addition to the target word identities). We used this information about HMM states to compute model RDMs for each of 40 British English phones in a procedure illustrated schematically in Fig 2. In order to account for the differences in recording characteristics and speaking styles between the training data used to build the ASR models and the stimulus words, a speaker adaptation method based on maximum a posteriori (MAP) criterion was used to adapt the mean values of the GMMs [34] and this resulted in a word accuracy of 91.6%.

**Figure 2:**
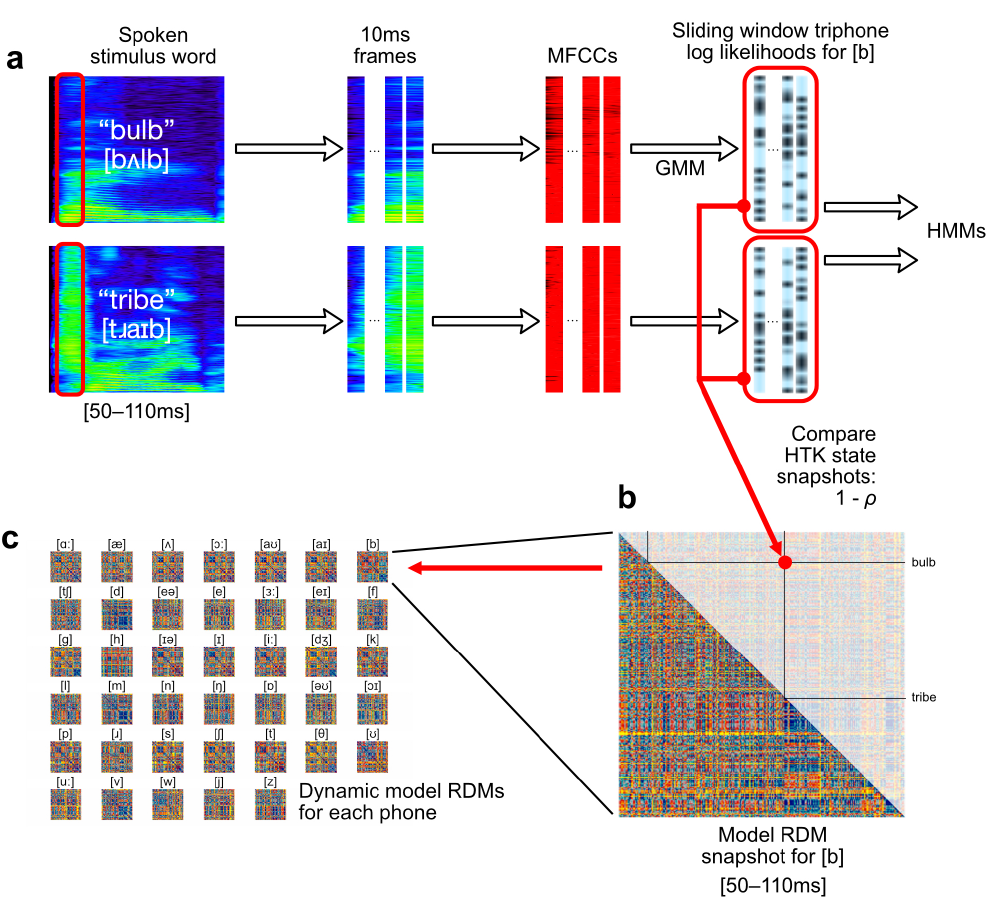
Mapping from GMM-HMM triphone log likelihoods to phone model RDMs. **(a)** Each 10 ms frame of audio is transformed into MFCC vectors. From these, a GMM estimates triphone log likelihoods, which are used in the phonetic HMMs. **(b)** We used the log likelihood estimates for each triphone variation of each phone, concatenated over a 60 ms sliding window, to model dissimilarities between input words. Dissimilarities modelled by correlation distances between triphone likelihood vectors were collected as entries in phonetic model RDMs. **(c)** These phone-specific model RDMs were computed through time for each sliding window position, yielding 40 time-varying model RDMs.

For performance reasons, the recorded stimuli were encoded into mel-frequency cepstral coefficients (MFCCs) [35] and represented each frame as a vector comprising 12 MFCCs plus the energy, together with their first and second derivatives, to give overall a 39dimensional real acoustic feature vector per frame [22]. Mel-frequency spectrogram transforms are similar to neurophysiologically motivated frequency transforms, such as Gammatone filterbank cochleagrams [6, 35, 36]. For each 10 ms frame, the GMM-based HMMs compute estimated log likelihoods that this frame was generated by particular hidden state sequence. A sketch of this process is shown in Fig 2a where the GMM-HMM triphone log likelihoods are used to compute model RDMs for each of our phones.

Specifically, for each phone *ϕ* present in the stimulus words, and since triphones correspond to contextual variations in pronunciation of phones, we grouped triphone log likelihood estimates by their triphone’s central phone, further concatenating the six 10-ms frames falling within each 60 ms sliding window. For each sliding window position and each phone *ϕ*, this gave us a vector of triphone log likelihood values with length equal to six times number of triphones containing *ϕ* in the central position. We used these values to build incremental RDMs modelling the information relevant to the representation or recognition of each phone *ϕ*, treating the vector of triphone log likelihood values as the *ϕ* model’s representation of the word for that 60 ms interval of time. We defined the distance between word pairs as the correlation distance (1 — Pearson’s correlation) between the triphone log likelihood vectors (Fig 2b), repeating this procedure at each 10 ms time-step up to the end of the 270 ms epoch to which these analyses were restricted. These procedures were repeated for each of 40 phones which were most commonly found in our 400 stimulus words, yielding 40 collections of phone model RDMs (Fig 2c), each indexed by the 21 positions of the sliding window, totalling 840 400 × 400-sized model RDM frames overall.

We describe these time-indexed collections of model RDMs as dynamic RDMs (dRDMs). We use the whole time-course of a dRDM to model the corresponding time-course of dRDMs computed from brain data.

### 2.3 Mapping phones to features

The use of phone labels in ASR systems corresponds to the usual target of converting speech to text, as well as the standard phonetic characterisation of speech sounds. However, phones themselves can be characterised by articulatory phonetic features, which correspond to the place and manner of articulation with which the phones are spoken. Given the substantial evidence from earlier research discussed in Section 1.2, the best candidate for the low-dimensional speech analysis solution adopted by the human system is likely to be a featural analysis of this type. A standard set of phonefeature correspondences, as used here, is summarised in Fig 3. By orienting the analysis in terms of features, we also reduce the dimensionality of our models, and simplify the interpretation by not requiring that very similar phones (e.g. [i] and [i]) be distinctly represented. This is especially relevant given that the GMM component of the ASR system, which produces triphone likelihoods, is not subject to any top-down processes which could guide identification of specific phones in a context wider than its input window. This also alleviates significant problems that emerged with the GLM mapping method used when this was conducted on a strictly phonetic rather than featural basis (see Section 2.5).

**Figure 3:**
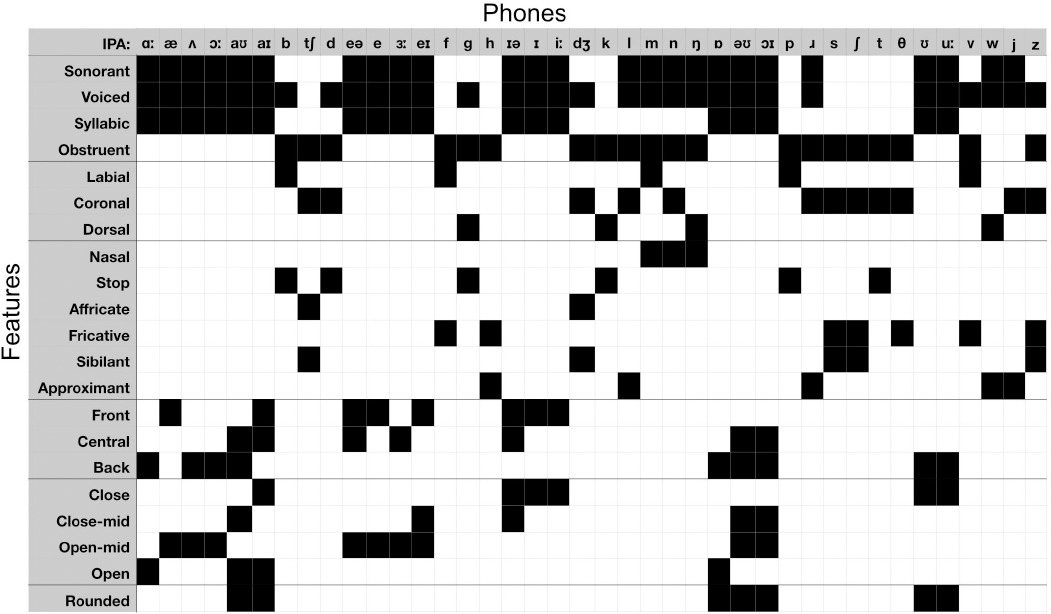
The mapping between articulatory features and phonetic labels. Columns describe phones and rows describe features, with a filled-in cell indicating that a phone exhibits a given feature. The features are grouped into descriptive categories, from top to bottom: Broad phonetic categories, place-of-articulation features for consonants, manner-of-articulation features for consonants, vowel frontness, vowel closeness, and vowel roundedness.

Since the choice of articulatory features rather than, for example, a more perceptually based feature set, is still open to discussion, and before committing ourselves to a reanalysis of the phone-based RDM data in these specific featural terms, we first evaluated the degree to which the chosen articulatory feature set was able to explain the second-order similarity structure of the set of phone model RDMs — i.e., the arrangement of each of the model RDMs in the same pairwise comparison space, described by the correlational structure of the model RDMs themselves [3, 5].

To conduct this novel analysis, we computed the pairwise Spearman correlation distances between the 79,800-length upper-triangular vectors of each pair of model RDMs at each timeframe, resulting in a 40 × 40-sized second-order model similarity matrix (Fig 4). The degree to which articulatory features explained the between-model similarity structure was evaluated by computing both Davies-Bouldin indices for the classification of models by each phonetic feature (Fig 5a), and *η*^2^ values representing the dissimilarity variance explained by the presence of each feature (Fig 5b). The Davies-Bouldin index is a standard way to evaluate the clustering of points in a high-dimensional space [37]. However, Davies-Bouldin index values cannot be judged in isolation, but only compared between different clusterings applied to the same data. An alternative is *η*^2^, values of which can be compared against fixed benchmarks [38].

**Figure 4:**
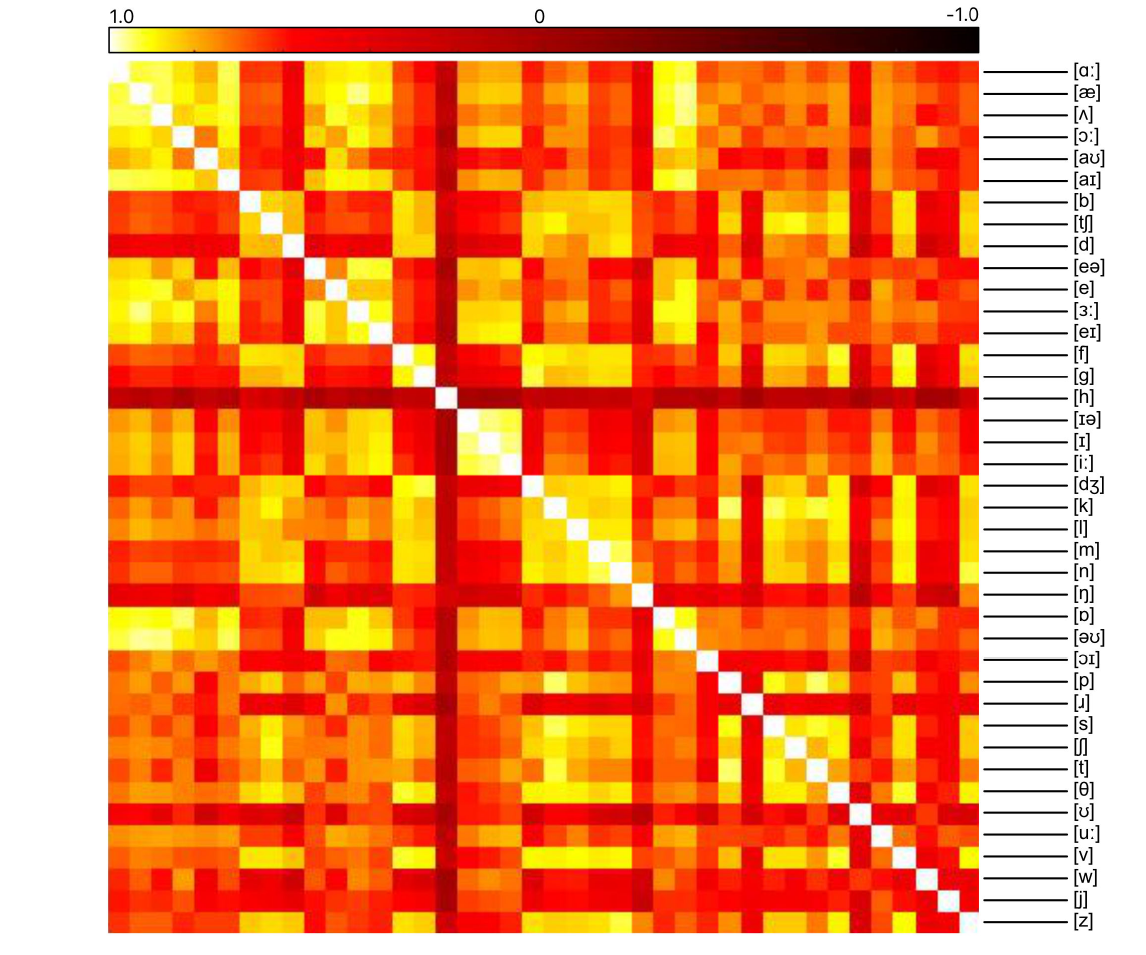
Second-order similarity structure of phone models. Entries in the matrix are Spearman’s rank correlations between model RDMs. The second-order similarity structure of the phone models for a representative time window centred over 90 ms after word onset, given by a correlation matrix between phone model RDMs.

**Figure 5:**
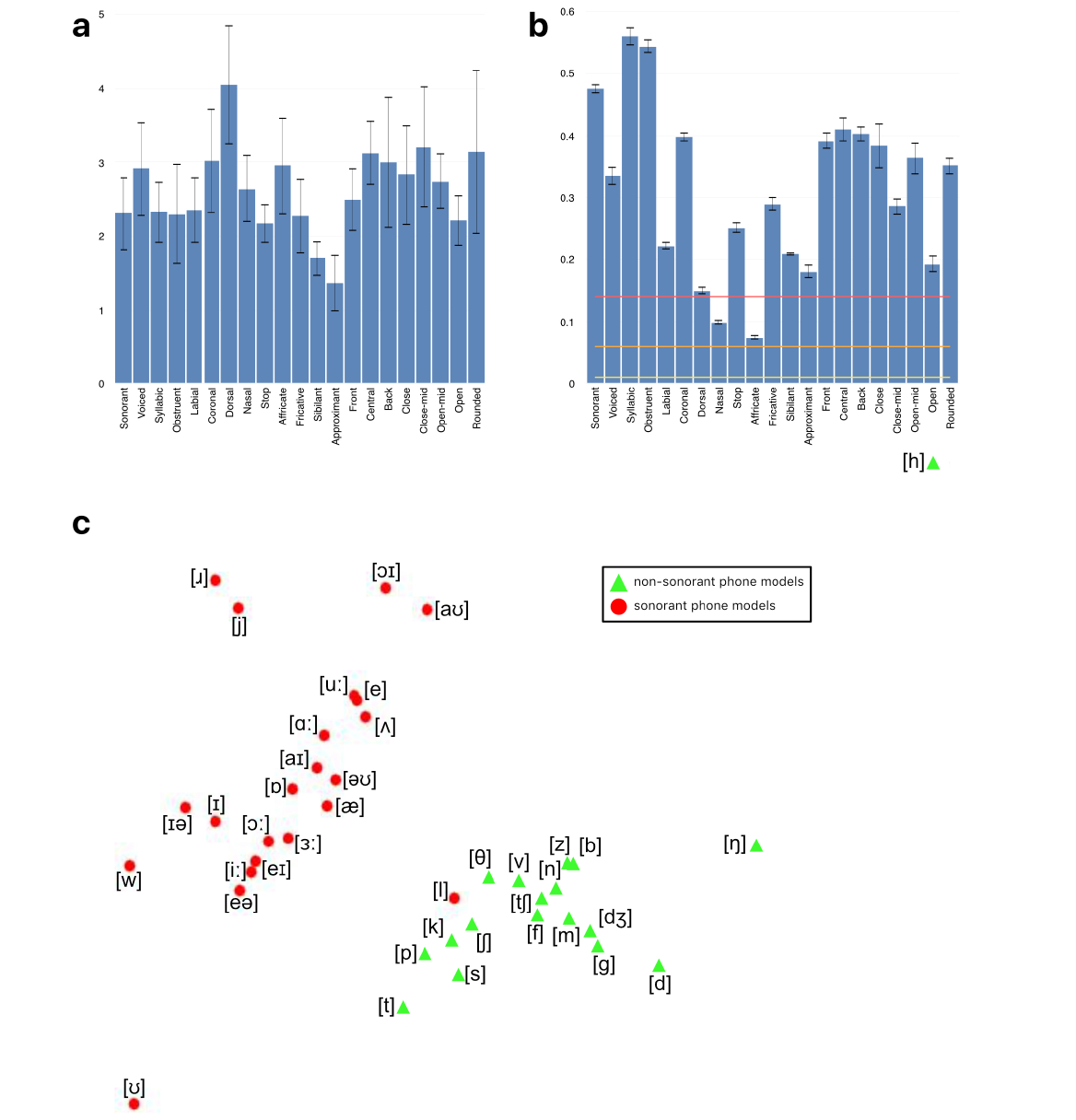
Similarities between model RDMs and phonetic features. **(a)** Davies-Bouldin indices for each feature. Error bars indicate one standard deviation of values over the epoch. **(b)** *η*^2^ values for each feature assignment. Rule-of-thumb guides for small, medium and large effect sizes are indicated by horizontal lines. Error bars indicate one standard deviation of values over the epoch. **(c)** The arrangement of phone models plotted using MDS (distance-dissimilarity correlation 0.94). Points are labelled according to the presence or absence of the sonorant feature. Models with the sonorant feature are represented with red circles, and models without the sonorant feature are represented with green triangles. The MDS arrangement of points as displayed was independent of the feature labelling.

We found that the arrangement of the phone models given by their similarity structure could be described in terms of the phonetic features possessed by the phones. As an example, we visualise this using a multidimensional scaling (MDS) plot [39] computed from the data at one particular timepoint. In this visualisation, each phone model is represented as a point in the plane (Fig 5c). The phone models appear to fall into two distinct classes, which are almost perfectly described by the presence of the sonorant feature. The relative distances between the models in the MDS plot are distorted from their true values by compression into the two dimensions of the plane, so that such a figure is useful for illustrative purposes only. However in this case the distortion is minimal (distance-dissimilarity correlation of 0.94). The majority of the articulatory features give large *η^2^* values, and their Davies-Bouldin indices are mostly comparable, showing that each feature contributes to the explanation of the models’ arrangement, and that no particular feature dominates the others (although sonorance and vowel place/position features tended to explain better than consonant manner features).

From this, it can be seen that the HTK GMM-HMM-derived model RDMs’ configuration are well explained by all phonetic features we used, providing statistically robust validation for the feature set adopted here.

## 2.4 Computing data dRDMs from incremental brain states

The second critical component of the RSA procedure is the computation of data RDMs (in this case brain data RDMs), whose similarity to the representational geometry of the model RDMs can then be determined. As the basis for these computations we used the existing EMEG source space data already collected for the 400 words entered into the GMM-HMM analysis described in Section 2.2. Sixteen right-handed native speakers of British English (six males, mean age 25 years, age range 19-35, self-reported normal hearing) were used from the original study. Recordings of 400 English words, as spoken by a female native British English speaker, were presented binaurally to participants. Each word was repeated once. The study was approved by the Peterborough and Fenland Ethical Committee (UK). For further details on data collection, see [32] and [1].

### 2.4.1 Source reconstruction

We estimated the location of cortical sources using the anatomically constrained minimum norm estimate (MNE) [24]. MR structural images for each participant were obtained using a GRAPPA 3D MPRAGE sequence (TR = 2250 ms; TE = 2.99 ms; flip-angle = 9deg; acceleration factor = 2) on a 3 T Trio (Siemens, Erlangen, Germany) with 1 mm isotropic voxels. From the MRI data, a representation of each participant’s cerebral cortex was constructed using FreeSurfer software (http://surfer.nmr.mgh.harvard.edu/). The forward model was calculated with a three-layer Boundary Element Model (BEM) using the outer surface of the scalp as well as the outer and inner surfaces of the skull identified in the anatomical MRI. This combination of MRI, MEG, and EEG data provides better source localization than MEG or EEG alone. The constructed cortical surface was decimated to yield approximately 12,000 vertices that were used as the locations of the dipoles. This was further restricted to the bilateral superior temporal mask as discussed previously. After applying the bilateral STC mask, 661 vertices remained in the left hemisphere and 613 in the right. To perform group analysis, the cortical surfaces of individual subjects were inflated and aligned using a spherical morphing technique implemented by MNE [25]. Sensitivity to neural sources was improved by calculating a noise covariance matrix based on the 100 ms pre-stimulus period. The activations at each location of the cortical surface were estimated over 1 ms windows.

### 2.4.2 Spatiotemporal searchlight RSA

The source-reconstructed representation of the electrophysiological activity of the brain was computed for participants’ responses to each of the target set of 400 words, and this was used to compute word-by-word RDMs in the following way: In ssRSA, we calculate RDMs from the EMEG data contained within a regular spatiotemporal searchlight patch of radius 20 mm and a 60 ms fixed-width sliding temporal window (see schematic diagram in Fig 6). This patch is moved to centre on each vertex in the source mesh, while the sliding window is moved throughout the epoch in fixed time-steps of 10 ms. From within each searchlight patch, we extracted the response pattern from each subject’s EMEG data from vertices within the patch and for time-points within the sliding window (Fig 6a). For each position of the sliding window, we computed a word-by-word RDM from these response patterns using the Pearson’s correlation distance measure of the resulting vectors (Fig 6b). We collected these into word-by-word dRDMs over the epoch. These dRDMs were averaged across subjects, resulting in one data RDM frame for each within-mask vertex and each sliding window position. This is repeated for the responses from each pair of experimental stimuli to produce a total set of 26,754 data RDMs, each associated with a specific vertex at a specific time-point.

**Figure 6:**
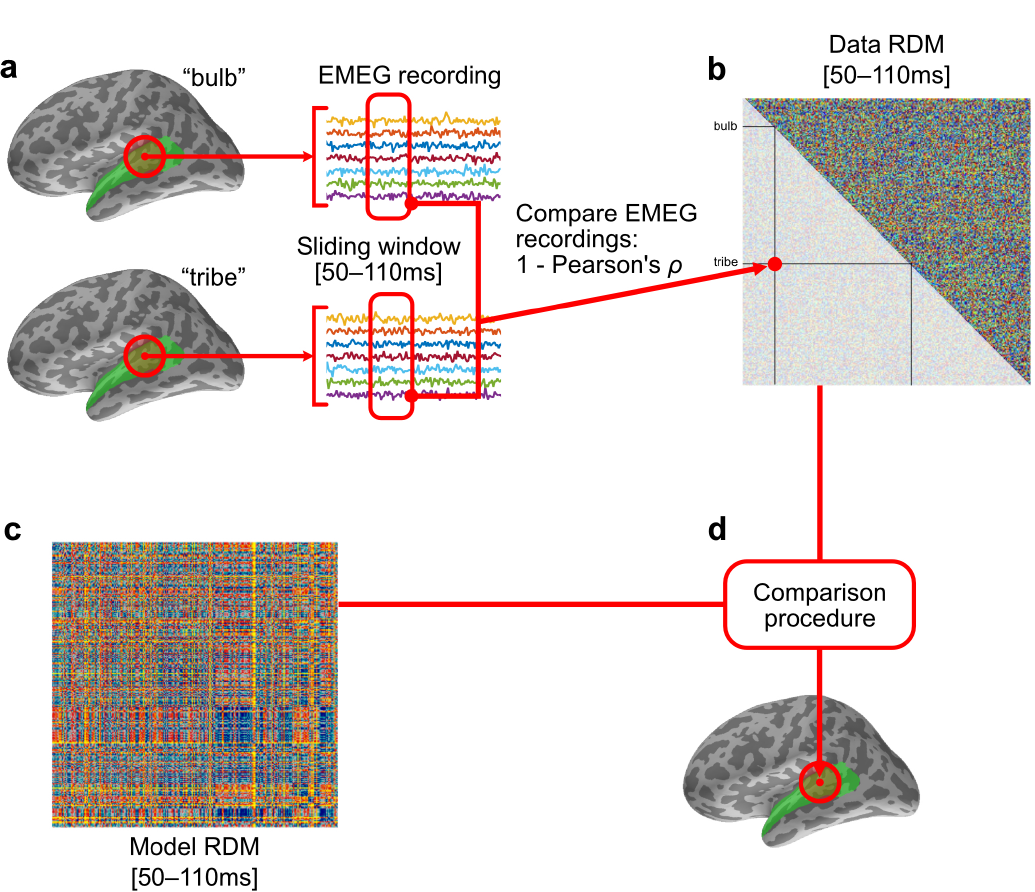
ssRSA for EMEG data. **(a)** Each stimulus’s evoked EMEG response is captured within a fixed time window and regular searchlight patch, which moves continuously in space inside the searchlight mask. **(b)** The dissimilarity between a pair of conditions is taken to be the correlation distance (1 — Pearson’s correlation) between the vectors of activation within the spatiotemporal searchlight. **(c)** The modelled dissimilarities between pairs of conditions are collated into a model RDM (see Fig 2). **(d)** Model RDMs are compared to the data RDM using a comparison procedure (see Section 2.5), with the resulting statistic mapped back into the central vertex of the searchlight. This is repeated for each spatiotemporal position of the searchlight.

## 2.5 Fitting model dRDMs to data dRDMs

To complete the RSA analysis process, each brain data RDM was compared to the 40 ASR-derived phone model RDMs computed for the relevant matching time-window, taking into account the 100 ms processing time-lag selected for these analyses (note that this displaces the neural analysis epoch to run from 100 to 370 ms from acoustic onset). Following the method of [1], multiple model RDMs can simultaneously be tested against data RDMs in the spatiotemporal searchlight using a generalized linear model (GLM). For a given data RDM D and corresponding 40 model RDMs *M*_[*a*]_,…, *M*_[*z*]_ for each phone, the GLM optimises coefficients *β*_1_, *β*_[*a*]_,…, *β*_[*z*]_ so as to minimise the sum-squared error *E* in the equation

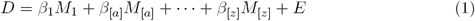

The *β*_*ϕ*_ coefficients then describe the contribution of each phone model *M*_*ϕ*_ to explaining the data, with *β*_1_ the coefficient for constant predictor *M*_1_ (Fig 7a). As the searchlight moved throughout time, we chose corresponding frames from the model dRDMs to use as predictors.

**Figure 7:**
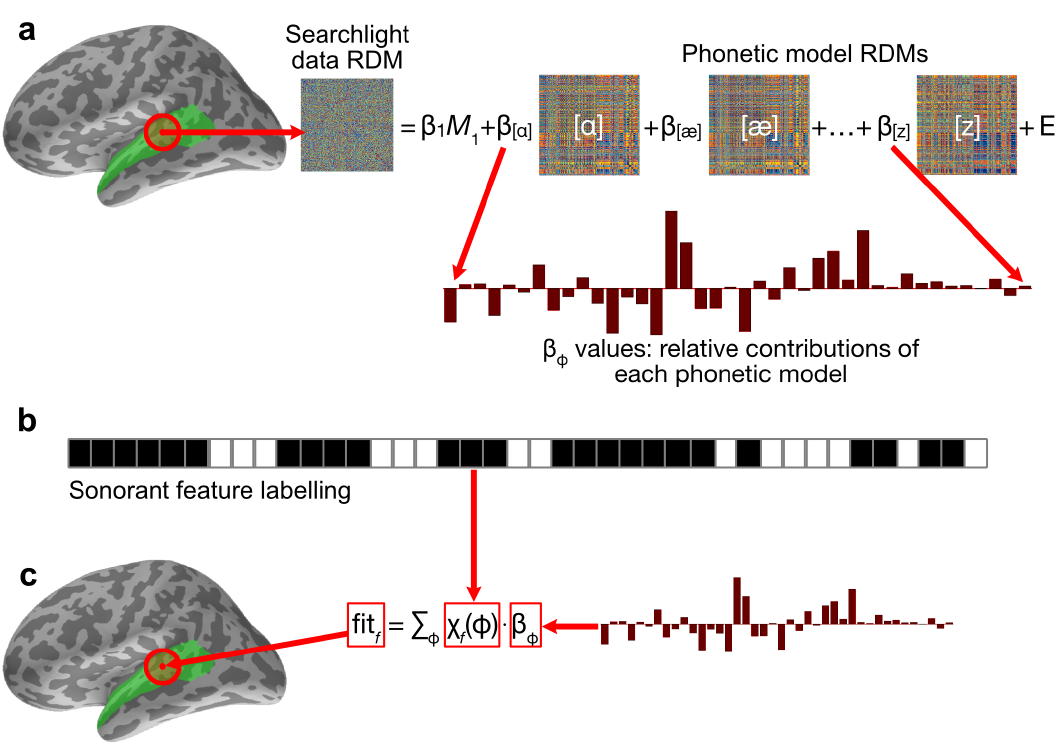
Relating brain data dRDMs to phone model dRDMs and converting to feature fits. **(a)** At each vertex and time point, all phone model RDMs are computed (Fig 6) and fitted against the data RDM in a GLM, yielding coefficients *β*_*ϕ*_. **(b)** The rows of the phone-feature matrix of Fig 3 describe for each feature *f* the phones *ϕ* exhibiting *f*, providing a labelling function *Xf*. The example given here is for the feature sonorant, the top row of the feature matrix in Fig 3. **(c)** The coefficients *β*_*ϕ*_ were aggregated by sum over each feature *f* to produce a map of fit for each feature, which was mapped back to the central location of the spatiotemporal searchlight.

In the second stage of the analysis procedure we convert phone model coefficients in each searchlight location GLM into their corresponding feature values (see Figs 7b and 7c). To do so we define the fit of a feature *f* at each spatiotemporal location to be the weighted sum

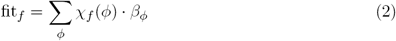

where *χ_f_* (*ϕ*) is 1 when phone *ϕ* has feature *f* and 0 otherwise (rows of feature matrix in Fig 3). Using this definition, we converted the phone coefficients *β*_*ϕ*_ into fit_*f*_ values describing the degree to which each feature *f* contributes to explaining the data (Fig 7b). These fit_*f*_ values can be mapped back into the centre of the searchlight patch from where the data RDM was calculated giving a spatiotemporal map of fit for each feature *f* (Fig 6d; Fig 7c). As well as being theoretically motivated (see Section 2.3), the aggregation of *β*_*ϕ*_ values over features was also preferred for practical reasons. HTK GMM-HMM models tended to assign similar log likelihoods to acoustically similar phones (e.g. [e] and [ea]), and as such the *β*_*ϕ*_ coefficients found in the GLM were affected by the presence of similar ‘competitor’ phones. Other phones which have no similar competitors (e.g. [k]) did not suffer from this. This source of systematic failure of the GLM analysis process meant that we were not able to compute successful incremental interpretations of the RSA model fitting process in phone-based terms. In contrast, by aggregating the *β*_*ϕ*_ coefficients according to features, we were able to link together the contributions of phone models with similar model dRDMs (Fig 5), and thereby provide a potentially more accurate account of the acoustic-phonetic information present in the signal. This advantage of a feature-based approach is consistent with suggestions in the behavioural and ASR literatures [10, 11, 40], pointing to the flexibility of such an approach in allowing partial acoustic-phonetic cues to be identified and exploited as they become incrementally available — contrasting with decisions about phone or speech sound identity, which may only become secure (if at all) when all of the relevant acoustic input as become available.

### 2.5.1 Statistical mapping

We computed statistical maps using a permutation method. Under the null hypothesis *H*_0_, there is no difference in the phonetic information between each word which would be represented in the brain responses, and thus we may randomly permute the condition labels for each of the words in our data RDMs, and would expect no difference in any fit of any model [1, 3, 41].

A null distribution of *β*_*ϕ*_ values was therefore simulated by randomly permuting the rows and columns of the data RDM [3], and re-computing the *β*_*ϕ*_ coefficients, and fit_*f*_ value. These were collected into separate null distributions of feature fits for each feature *f*, pooling the values over the vertices within the searchlight mask. Separate null distributions were pooled for each feature as different numbers of phones contribute to each feature. This provides a permutation-derived simulated null distributions of more than 10,000 samples for each feature.

By taking a certain quantile (e.g. 0.95) for each of these null fit_*f*_ distributions, we obtain a confidence threshold *θ_f_* for each feature *f*. We use *θ_f_* to threshold each fit_*f*_-map. For the purposes of providing a summary of the whole time epoch, we averaged the *β*_*ϕ*_ values over the 100-370 ms epoch (see Section 1.3) before fitting features in both the individual maps and the calculation of the θ_*f*_ thresholds.

## 2.6 RSA evaluation of ASR models in neural source space

Fig 8a shows each of the thresholded feature maps in ensemble. Where a feature showed significant fit for any vertex, the most conservative possible standard *p*-threshold (*p* < 0.05, 0.01 or 0.001) was chosen for which the cluster remained. This gave the fullest picture of the results. The maps are somewhat different between hemispheres, though clusters appear over bilateral Heschl’s gyri, particularly concentrated in primary auditory regions such as A1. Both hemispheres show a wide regions of fit, parcellated into partially overlapping patches for different features. Most of the features we tested showed significant fit. The exceptions were the features affricate, fricative and approximant, for which no vertices within the searchlight mask showed significant fit.

**Figure 8:**
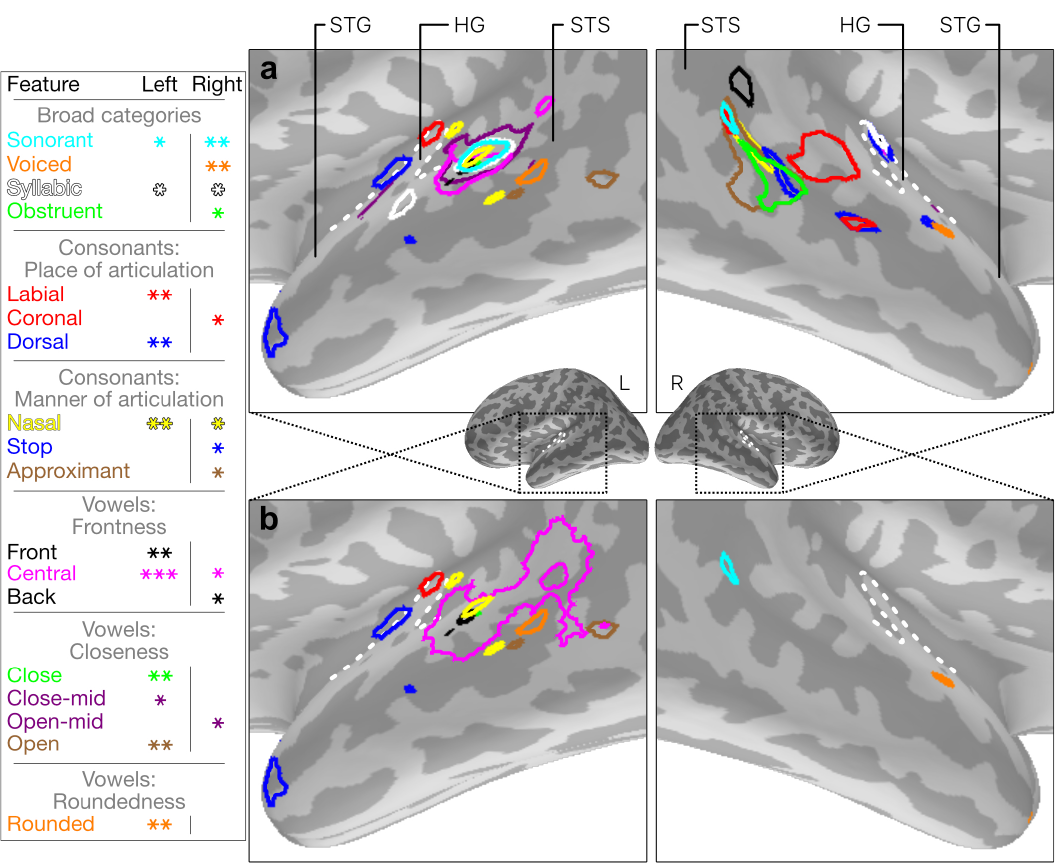
Maps of fit for each feature. **(a)** Thresholded maps of fit for each feature for which at least one vertex showed significant fit between 100 ms and 370 ms. We report *p* < 0.05 using *, *p* < 0.01 using **, and *p* < 0.001 using ***. Light and dark greys represent gyri and sulci respectively. The miniature figures show the location of the larger diagrams. Anatomical landmarks are superior temporal gyrus (STG), superior temporal sulcus (STS) and Heschl’s gyrus (HG). The dotted white lines indicate the outline of HG. **(b)** With a fixed threshold of *p* < 0.01 for both hemispheres, only broad-category features remain on the right, and within-category distinctive features are dominant on the left

Features describing broad phonetic category distinctions (sonorant, voiced and obstruent) all showed greater fit within the right hemisphere. Feature fits in the left hemisphere tended to be more focussed than in the right and more significant (indicated by asterisks on the figure). Features which showed fits in both hemispheres were those providing finer within-category distinctions, such as central and nasal. To visualise this further, we fixed a threshold at *p* < 0.01 for both hemispheres and all feature maps (see Fig 8b). This left intact some broad category matches in right STG, and suggested a greater coverage in left STG, STS and HG for within-category feature distinctions — place and manner of articulation features for consonants, and closeness, frontness and roundedness dimensions for vowels.

## 3 Discussion

In this study we have used a functional model of machine speech recognition to investigate early representations of speech in human superior temporal cortex (STC). By using RSA, we have been able to directly compare representations of speech in the machine system with representations found in EMEG activations measured from human participants. We mapped the space of phones used by the ASR to a smaller space of articulatory features, after verifying that these features acceptably describe the similarity structure of the ASR representations. We tested feature fits systematically throughout the STC regions of each of our participants, using a spatiotemporal searchlight technique, and looked at vertices showing consistent fit across our participants. A vertex shown to be significant in our results (Fig 8) indicates the location of spatially local patterns of activation for each of our stimulus words which is explained by information relevant to a particular feature, as modelled by the ASR system’s representation. We found distributed patches of sensitivity to each articulatory feature throughout bilateral STC. The distribution exhibited some left-right asymmetry, with broad category features (e.g. sonorant) fitting best in the left hemisphere, and more fine-grained within-category features fiting better in the right. Overall, fit in the left hemisphere was more significant.

These results confirm the feasibility of combining ssRSA with source space EMEG to relate dynamic brain states to dynamic machine states in the incremental speech input environment. In doing so, we see significant correspondences between human and ASR solutions to the speech recognition problems, captured here in terms of a low-dimensional representational framework related to articulatory phonetic features. We discuss below the empirical status of these findings, followed by a more general analysis of the meaning of potential brain-machine correspondences in the speech recognition environment.

### 3.1 Phonetic feature sensitivity in human temporal cortex

The critical result in this study is that phonetic model RDMs, extracted from the ASR labelling of the phonetic content of incremental 60 ms speech input time windows, could be significantly related to brain data RDMs capturing the human neural response in superior temporal cortex to the same incremental time windows. These relationships took the form of spatially coherent patches in STC whose responses showed significant fit to a variety of different articulatory phonetic features. Is there independent empirical evidence that bears on the credibility of this finding of distributed patches of featural sensitivity in STC?

The overall result — supporting the viability of an account in terms of features — is compatible with many strands of evidence favouring such an approach to the neural representation of speech in human temporal cortex. Most directly, it is fully consistent with the Mesgarani et al. [19] results, showing that ECoG electrode responses in STC can be well characterised in terms of specific phonetic features. Given the isomorphism, however, between featural and non-featural phonetic representations, it is important to be clear that neither Mesgarani’s results nor our own can by themselves exclude accounts of neural coding in STC based on a classic phone-based approach. Equally, it should be clear that the current results, while consistent with a featural decomposition of ASR representations of regularities in the speech-to-phone mapping, cannot exclude a variety of other accounts, including those built entirely from acoustically defined regularities.

The second salient aspect of the results is the computational geography that they reveal, with different cortical patches showing a preference for specific features, and where these patches are scattered in partially overlapping fashion in auditory cortex and in surrounding posterior STG and STS. Although there is currently no neurocomputationally or neuroanatomically explicit account of how STC supports acoustic-phonetic analysis, the geography of these results is broadly consistent with existing data from other neuroimaging studies. The regions of superior temporal and Heschl’s gyri which showed significant fit for at least one phonetic feature in our results are consistent with areas which have shown speech selectivity in other studies [12–15, 19, 42, 43]. In addition, the spatial locations of these featural patches overlap with regions found to have frequency-selective responses to both high and low frequencies, and both wide and narrow frequency tuning curves [1, 43–46]. Turning to more specific studies of the neural representations of phonological features during speech perception, the Arsenault and Buchsbaum fMRI study [47] shows bilateral regions of superior temporal cortex supporting discrimination between articulatory feature categories. Closest to our current results, the fMRI study by Correia et al. [48], using a searchlight MVPA technique, not only showed that brain responses to syllables differing in their voicing, place or manner of articulation could be distinguished in STC, but also that they have somewhat patchy areas of distribution. Finally, the older study by Obleser et al. [14], using MVPA techniques on an fMRI data set, provides similar evidence for a distribution of vowel and consonant feature-sensitive patches across human STC.

While these resemblances between studies are not sufficient to validate the specific feature-sensitive regions found in the present study, they do suggest that the computational geography that our results indicate is compatible with current independent findings of regions in STC sensitive to specific featural dimensions of the speech input. They are also compatible with evidence from other domains that the style of computation found for complex species-specific processes can have similar patch-like properties. Research by Grimaldi et al. [49], for example, reveals a network of different face patches in the macaque (as well as in humans), which are densely interconnected in both feedforward and feedback directions to constitute a highly specialised processing network. Given the evolutionary significance of human speech recognition, similar networks may underpin the even more specialised and species-specific process of recovering the articulatory gestures underlying a given speech input.

### 3.2 Relating machine states to brain states: vision and speech

On the assumption that there is empirical support for the outcomes reported here, we now turn to the question of what it might mean to find correspondences between machine-based solutions to a complex mapping and the ‘natural’ neural solutions found in the brains of humans or other species. It is important to distinguish here between the visual and the speech domains. There is a sizeable and rapidly growing body of work which relates computational models in the domains of visual object classification and visual scene analysis to the properties of the corresponding neural processes in humans and other primates (e.g. [26, 50–53]). This is not the case for the speech domain.

The work on vision differs from speech and language research in several major respects. The first is that the neural processing machinery that supports visual perceptual processing is essentially the same, in its major respects, in humans and in other primates. There is extensive evidence (e.g. [50, 54–57]) for similarities both in the basic architecture of the system — the primate dorsal and ventral visual processing streams — and in the computational properties of the analyses carried out, based on a complex multi-level hierarchical cortical organisation. This means that, unlike speech comprehension, there are detailed models of the sequence of processes that map from early visual cortex to higher-order perceptual and conceptual representations in more anterior brain regions.

Second, and closely related to the first, there is a strong tradition of computational implementations of models of visual perceptual processing that reflect the theories of hierarchical cortical organisation proposed for different aspects of these processes, and where these design assumptions (such as localised convolution and pooling) are also reflected in machine vision implementations. These relationships have been greatly strengthened by the recent emergence of deep neural network (DNN) systems as the machine learning systems of choice, and where several studies suggest parallels between the representations computed in the successive layers of such DNNs and the successive levels hypothesized for the human system [50, 53]. Importantly, this means that the internal structures of machine vision systems are potentially informative and relevant to our understanding of the neurocomputational architecture of the natural system (and *vice versa*), and not just whether they generate equivalent outputs (for example in object classification tasks).

None of these factors currently hold true for speech comprehension, either by humans or by machines. While the human auditory processing system does have close parallels with the general primate system [29], no non-human primate supports anything like the human system of speech communication, where intricately modulated sequences of speech sounds, uttered by members of the same species, map onto tens of thousands of learned linguistic elements (words and morphemes), each with its own combination of acoustic-phonetic identifiers. There is simply no primate model of how these mappings are achieved (cf. [58]). No doubt partly because of this, but also reflecting the complexities of the real-time analyses the human brain carries out on an incremental and dynamically varying sensory input, there are no computationally explicit neurocognitive models of how humans solve the spoken word-recognition problem.

Consistent with this, and following design strategies long established in the speech engineering domain (cf. [59]), ASR systems have been driven (with considerable success) solely by computational engineering considerations, such as statistical learning efficiency, with little or no reference to the properties of the human system. It was not a goal in the design and implementation of such systems that they should correspond to the supposed architecture of the human system, nor have there been any compelling demonstrations that performance would benefit if such considerations were taken into account. This holds true not only for the kind of GMM-HMM architecture used here, but also for the DNN-based systems that have recently emerged [60, 61].

In the current research, therefore, the focus has been (as noted earlier) on the input-output relations exhibited by the two types of system — human superior temporal cortex and automatic speech recogniser — with ssRSA allowing us to abstract away from the very different formats of representations in each of these systems. Strikingly, as documented in this paper, we have been able to show that the regularities that successful ASR systems encode in the mapping between speech input and word-level phonetic labelling can indeed be related to the regularities extracted by the human system as it processes and identifies parallel sets of words. What are implications of this result?

### 3.3 Implications and conclusions

The finding of significant machine-brain correspondences suggests, first, that there are major underlying commonalities in the solutions of human and machine systems to the speech recognition problem. In particular, the combination of ssRSA and EMEG data sets, as implemented here, suggest that such relations can be informed and guided by direct inspection and comparison of dynamic machine states and dynamic EMEG-derived brain states, adding a novel empirical dimension to such comparisons.

Second, the finding that the commonalities between human and machine systems can be characterised in terms of a biologically plausible low-dimensional analysis, based on articulatory phonetic features, argues for an increased focus on the neurobiological and computational substrate for such an analysis strategy. In ASR research, the development of systems based around the extraction of articulatory features has a long history (e.g., [11]) and some recent exemplars (e.g., [62]), but is not central to most current work. It would greatly strengthen future investigations of potential links between proposed human feature-based analysis processes and the representational strategies in machine recognition systems if ASR systems trained to extract featural representations could be tested against human brain data in the same way as the phone-based system tested here. In the human domain, the evidence for specific phonetic feature representations in superior temporal cortex raises several further questions about the nature of the fine-grained neural representations within these patches, and how these might relate to the neural reconstruction of underlying featural representations.

Third, and finally, the successful matching of the correlational structure of input-output solutions in human and machine systems may help to motivate the development of more biologically informed approaches to ASR. Such approaches are now commonplace, for example, in machine vision systems as discussed previously, and may prove equally valuable in future ASR systems. This will prove an equally positive development for the study of the human system, which stills lacks explicit neurocomputational “end-to-end” models of the process of understanding speech.

## Acknowledgements

The authors thank Niko Kriegeskorte, Lorraine K. Tyler and Barry Devereux for thoughtful comments and discussions. RSA computation was done in Matlab using the open-source RSA toolbox [3] and custom EMEG extensions, to which Isma Zulfiqar, Fawad Jamshed and Jana Klímová also contributed code. Davies-Bouldin computations were performed using the SOM toolbox for Matlab 5 [63].

This research was supported financially by an Advanced Investigator grant to WMW from the European Research Council (AdG 230570 NEUROLEX), by MRC Cognition and Brain Sciences Unit (CBSU) funding to WMW (U.1055.04.002.00001.01), and by a European Research Council Advanced Investigator grant under the European Communitys Horizon 2020 Research and Innovation Programme (2014-2020 ERC Grant agreement no 669820) to Lorraine K. Tyler. LS was partly supported by the NIHR Biomedical Research Centre and Biomedical Unit in Dementia based at Cambridge University Hospital NHS Foundation Trust.

## Author contributions

CW and LS conceived of analyses and wrote analysis scripts. CW conducted the analyses. XL, CZ and PW contributed ASR software, technical assistance and valuable advice. EF designed, conducted and analysed the EMEG spoken words experiment. AT preprocessed the neuroimaging data. WMW designed the experiment, conceived of the analyses and supervised the project. CW and WMW wrote the manuscript. All authors discussed the results and implications and commented on the manuscript at all stages.

## References

1 L. Su, I. Zulfiqar, F. Jamshed, E. Fonteneau, and W. Marslen-Wilson, “Mapping tonotopic organization in human temporal cortex: representational similarity analysis in EMEG source space,” Frontiers in neuroscience, vol. 8, 2014.

2 N. Kriegeskorte, R. Goebel, and P. Bandettini, “Information-based functional brain mapping,” Proceedings of the National Academy of Sciences of the United States of America, vol. 103, no. 10, pp. 3863–3868, 2006.

3 H. Nili, C. Wingfield, A. Walther, L. Su, W. Marslen-Wilson, and N. Kriegeskorte, “A toolbox for representational similarity analysis,” PLoS Comput. Biol, vol. 10, no. 4, p. e1003553, 2014.

4 L. Su, E. Fonteneau, W. Marslen-Wilson, and N. Kriegeskorte, “Spatiotemporal searchlight representational similarity analysis in EMEG source space,” in Pattern Recognition in NeuroImaging (PRNI), 2012 International Workshop on, pp. 97–100, IEEE, 2012.

5 N. Kriegeskorte, M. Mur, and P. Bandettini, “Representational similarity analysis-connecting the branches of systems neuroscience,” Frontiers in systems neuroscience, vol. 2, 2008.

6 R. D. Patterson, “A pulse ribbon model of monaural phase perception,” The Journal of the Acoustical Society of America, vol. 82, no. 5, pp. 1560–1586, 1987.

7 N. Kriegeskorte and R. A. Kievit, “Representational geometry: integrating cognition, computation, and the brain,” Trends in cognitive sciences, vol. 17, no. 8, pp. 401–412, 2013.

8 J. L. Elman, “On the meaning of words and dinosaur bones: Lexical knowledge without a lexicon,” Cognitive Science, vol. 33, no. 4, pp. 547–582, 2009.

9 W. Marslen-Wilson and P. Warren, “Levels of perceptual representation and process in lexical access: words, phonemes, and features.,” Psychological review, vol. 101, no. 4, p. 653, 1994.

10 P. Warren and W. Marslen-Wilson, “Continuous uptake of acoustic cues in spoken word recognition,” Perception & Psychophysics, vol. 41, no. 3, pp. 262–275, 1987.

11 L. Deng and D. X. Sun, “A statistical approach to automatic speech recognition using the atomic speech units constructed from overlapping articulatory features,” The Journal of the Acoustical Society of America, vol. 95, no. 5, pp. 2702–2719, 1994.

12 J. Obleser, A. Lahiri, and C. Eulitz, “Magnetic brain response mirrors extraction of phonological features from spoken vowels,” Journal of Cognitive Neuroscience, vol. 16, no. 1, pp. 31–39, 2004.

13 J. Obleser, H. Boecker, A. Drzezga, B. Haslinger, A. Hennenlotter, M. Roettinger, C. Eulitz, and J. P. Rauschecker, “Vowel sound extraction in anterior superior temporal cortex,” Human brain mapping, vol. 27, no. 7, pp. 562–571, 2006.

14 J. Obleser, A. Leaver, J. VanMeter, and J. P. Rauschecker, “Segregation of vowels and consonants in human auditory cortex: evidence for distributed hierarchical organization,” Frontiers in psychology, vol. 1, p. 232, 2010.

15 M. Scharinger, U. Domahs, E. Klein, and F. Domahs, “Mental representations of vowel features asymmetrically modulate activity in superior temporal sulcus,” Brain and Language, vol. 163, pp. 42–49, 2016.

16 R. Möttönen and K. E. Watkins, “Motor representations of articulators contribute to categorical perception of speech sounds,” The Journal of Neuroscience, vol. 29, no. 31, pp. 9819–9825, 2009.

17 A. D. Ausilio, F. Pulvermöller, P. Salmas, I. Bufalari, C. Begliomini, and L. Fadiga, “The motor somatotopy of speech perception,” Current Biology, vol. 19, no. 5, pp. 381–385, 2009.

18 F. Pulvermüller, M. Huss, F. Kherif, F. M. del Prado Martin, O. Hauk, and Y. Shtyrov, “Motor cortex maps articulatory features of speech sounds,” Proceedings of the National Academy of Sciences, vol. 103, no. 20, pp. 7865–7870, 2006.

19 N. Mesgarani, C. Cheung, K. Johnson, and E. F. Chang, “Phonetic feature encoding in human superior temporal gyrus,” Science, vol. 343, no. 6174, pp. 1006–1010, 2014.

20 P. Ladefoged and K. Johnson, A course in phonetics. Cengage Learning, Scarborough, 2011.

21 P. Roach, “British English: Received pronunciation,” Journal of the International Phonetic Association, vol. 34, no. 02, pp. 239–245, 2004.

22 S. Young, G. Evermann, D. Kershaw, G. Moore, J. Odell, D. Ollason, V. Valtchev, and P. Woodland, The HTK book (for HTK version 3.4.1). Cambridge University Engineering Department, 2009.

23 A. Molins, S. M. Stufflebeam, E. N. Brown, and M. S. Hömöläinen, “Quantification of the benefit from integrating meg and eeg data in minimum *l*_2_-norm estimation,” Neuroimage, vol. 42, no. 3, pp. 1069–1077, 2008.

24 M.S. Hamalainen and R. Ilmoniemi, “Interpreting magnetic fields of the brain: minimum norm estimates,” Medical & biological engineering & computing, vol. 32, no. 1, pp. 35–42, 1994.

25 A. Gramfort, M. Luessi, E. Larson, D. A. Engemann, D. Strohmeier, C. Brodbeck, L. Parkkonen, and M. S. Hamalainen, “MNE software for processing MEG and EEG data,” Neuroimage, vol. 86, pp. 446–460, 2014.

26 R. M. Cichy, A. Khosla, D. Pantazis, and A. Oliva, “Dynamics of scene representations in the human brain revealed by magnetoencephalography and deep neural networks,” NeuroImage, 2016.

27 E. F. Chang, J. W. Rieger, K. Johnson, M. S. Berger, N. M. Barbaro, and R. T. Knight, “Categorical speech representation in human superior temporal gyrus,” Nature neuroscience, vol. 13, no. 11, pp. 1428–1432, 2010.

28 M. H. Davis and I. S. Johnsrude, “Hierarchical processing in spoken language comprehension,” The Journal of Neuroscience, vol. 23, no. 8, pp. 3423–3431, 2003.

29 J. P. Rauschecker and S. K. Scott, “Maps and streams in the auditory cortex: nonhuman primates illuminate human speech processing,” Nature neuroscience, vol. 12, no. 6, pp. 718–724, 2009.

30 A. Thwaites, I. Nimmo-Smith, E. Fonteneau, R. D. Patterson, P. Buttery, and W. D. Marslen-Wilson, “Tracking cortical entrainment in neural activity: auditory processes in human temporal cortex,” Frontiers in computational neuroscience, vol. 9, 2015.

31 S. G. Wardle, N. Kriegeskorte, T. Grootswagers, S.-M. Khaligh-Razavi, and T. A. Carlson, “Perceptual similarity of visual patterns predicts the similarity of their dynamic neural activation patterns measured with meg,” arXiv preprint arXiv:1506.02208, 2015.

32 E. Fonteneau, M. Bozic, and W. D. Marslen-Wilson, “Brain network connectivity during language comprehension: Interacting linguistic and perceptual subsystems,” Cerebral Cortex, p. bhu283, 2014.

33 S. J. Young, J. J. Odell, and P. C. Woodland, “Tree-based state tying for high accuracy acoustic modelling,” in Proceedings of the workshop on Human Language Technology, pp. 307–312, Association for Computational Linguistics, 1994.

34 J.-L. Gauvain and C.-H. Lee, “Maximum a posteriori estimation for multivariate gaussian mixture observations of markov chains,” IEEE transactions on speech and audio processing, vol. 2, no. 2, pp. 291–298, 1994.

35 S. B. Davis and P. Mermelstein, “Comparison of parametric representations for monosyllabic word recognition in continuously spoken sentences,” Acoustics, Speech and Signal Processing, IEEE Transactions on, vol. 28, no. 4, pp. 357–366, 1980.

36 M. Pitz and H. Ney, “Vocal tract normalization equals linear transformation in cepstral space,” Speech and Audio Processing, IEEE Transactions on, vol. 13, no. 5, pp. 930–944, 2005.

37 D. L. Davies and D. W. Bouldin, “A cluster separation measure,” IEEE transactions on pattern analysis and machine intelligence, no. 2, pp. 224–227, 1979.

38 J. Cohen, “Statistical power analysis for the behavioural sciences”. hillside, NJ: Lawrence Earlbaum Associates, 1988.

39 R. N. Shepard, “Multidimensional scaling, tree-fitting, and clustering,” Science, vol. 210, no. 4468, pp. 390–398, 1980.

40 P. Warren and W. Marslen-Wilson, “Cues to lexical choice: Discriminating place and voice,” Perception & Psychophysics, vol. 43, no. 1, pp. 21–30, 1988.

41 T. E. Nichols and A. P. Holmes, “Nonparametric permutation tests for functional neuroimaging: a primer with examples,” Human brain mapping, vol. 15, no. 1, pp. 1–25, 2002.

42 A. M. Chan, A. R. Dykstra, V. Jayaram, M. K. Leonard, K. E. Travis, B. Gygi, J. M. Baker, E. Eskandar, L. R. Hochberg, E. Halgren, et al., “Speech-specific tuning of neurons in human superior temporal gyrus,” Cerebral Cortex, vol. 24, no. 10, pp. 2679–2693, 2014.

43 M. Moerel, F. De Martino, and E. Formisano, “An anatomical and functional topography of human auditory cortical areas,” Frontiers in neuroscience, vol. 8, 2014.

44 S. Baumann, C. I. Petkov, and T. D. Griffiths, “A unified framework for the organization of the primate auditory cortex,” Frontiers in systems neuroscience, vol. 7, 2013.

45 M. Saenz and D. R. Langers, “Tonotopic mapping of human auditory cortex,” Hearing research, vol. 307, pp. 42–52, 2014.

46 S. Norman-Haignere, N. Kanwisher, and J. H. McDermott, “Cortical pitch regions in humans respond primarily to resolved harmonics and are located in specific tonotopic regions of anterior auditory cortex,” The Journal of Neuroscience, vol. 33, no. 50, pp. 19451–19469, 2013.

47 J. S. Arsenault and B. R. Buchsbaum, “Distributed neural representations of phonological features during speech perception,” The Journal of Neuroscience, vol. 35, no. 2, pp. 634–642, 2015.

48 J. M. Correia, B. M. Jansma, and M. Bonte, “Decoding articulatory features from fmri responses in dorsal speech regions,” The Journal of Neuroscience, vol. 35, no. 45, pp. 15015–15025, 2015.

49 P. Grimaldi, K. S. Saleem, and D. Tsao, “Anatomical connections of the functionally defined face patches in the macaque monkey,” Neuron, 2016.

50 S.-M. Khaligh-Razavi and N. Kriegeskorte, “Deep supervised, but not unsupervised, models may explain it cortical representation,” PLoS Comput Biol, vol. 10, no. 11, p. e1003915, 2014.

51 N. Kriegeskorte, M. Mur, D. A. Ruff, R. Kiani, J. Bodurka, H. Esteky, K. Tanaka, and P. A. Bandettini, “Matching categorical object representations in inferior temporal cortex of man and monkey,” Neuron, vol. 60, no. 6, pp. 1126–1141, 2008.

52 A. Clarke, B. J. Devereux, B. Randall, and L. K. Tyler, “Predicting the time course of individual objects with meg,” Cerebral Cortex, p. bhu203, 2014.

53 U. Güclu and M. A. van Gerven, “Deep neural networks reveal a gradient in the complexity of neural representations across the ventral stream,” The Journal of Neuroscience, vol. 35, no. 27, pp. 10005–10014, 2015.

54 K. Denys, W. Vanduffel, D. Fize, K. Nelissen, H. Peuskens, D. Van Essen, and G. A. Orban, “The processing of visual shape in the cerebral cortex of human and nonhuman primates: a functional magnetic resonance imaging study,” The Journal of Neuroscience, vol. 24, no. 10, pp. 2551–2565, 2004.

55 G. A. Orban, D. Van Essen, and W. Vanduffel, “Comparative mapping of higher visual areas in monkeys and humans,” Trends in cognitive sciences, vol. 8, no. 7, pp. 315–324, 2004.

56 R. B. Tootell, D. Tsao, and W. Vanduffel, “Neuroimaging weighs in: humans meet macaques in “primate” visual cortex,” The Journal of Neuroscience, vol. 23, no. 10, pp. 3981–3989, 2003.

57 D. C. Van Essen, J. W. Lewis, H. A. Drury, N. Hadjikhani, R. B. Tootell, M. Bakircioglu, and M. I. Miller, “Mapping visual cortex in monkeys and humans using surface-based atlases,” Vision research, vol. 41, no. 10, pp. 1359–1378, 2001.

58 A. Amador and D. Margoliash, “A mechanism for frequency modulation in songbirds shared with humans,” The Journal of Neuroscience, vol. 33, no. 27, pp. 11136–11144, 2013.

59 F. Jelinek, Statistical methods for speech recognition. MIT press, 1997.

60 S. Young, G. Evermann, M. Gales, T. Hain, D. Kershaw, X. Liu, G. Moore, J. Odell, D. Ollason, D. Povey, A. R. V. Valtchev, P. Woodland, and C. Zhang, The HTK book (for HTK version 3.5). Cambridge University Engineering Department, 2015.

61 C. Zhang and P. C. Woodland, “A general artificial neural network extension for htk,” Proc. Interspeech, Dresden, 2015.

62 V. Mitra, W. Wang, A. Stolcke, H. Nam, C. Richey, J. Yuan, and M. Liberman, “Articulatory trajectories for large-vocabulary speech recognition,” in 2013 IEEE International Conference on Acoustics, Speech and Signal Processing, pp. 7145–7149, IEEE, 2013.

63 J. Vesanto, J. Himberg, E. Alhoniemi, and J. Parhankangas, “Som toolbox for matlab 5,” in Technical Report A57, Helsinki University of Technology, 2000.

